# Partial correlation analysis of transcriptomes helps detangle the growth and defense network in spruce

**DOI:** 10.1101/247981

**Authors:** Ilga Porth, Richard White, Barry Jaquish, Kermit Ritland

## Abstract

- In plants, there can be a trade-off between resource allocations to growth versus defense. Here, we use partial correlation analysis of gene expression to make inferences about the nature of this interaction.
- We studied segregating progenies of Interior spruce subject to weevil attack. In a controlled experiment, we measured pre-attack plant growth and post-attack damage with several morphological measures, and profiled transcriptomes of 188 progeny.
- We used partial correlations of individual transcripts (ESTs) with pairs of growth/defense traits to identify important nodes and edges in the inferred underlying gene network, e.g., those pairs of growth/defense traits with high mutual correlation with a single EST transcript. We give a method to identify such ESTs.
- A terpenoid ABC transporter gene showed strongest correlations (*P*=0.019); its transcript represented a hub within the compact 166-member gene-gene interaction network (*P*=0.004) of the *negative* genetic correlations between growth and subsequent pest attack. A small 21-member interaction network (*P*=0.004) represented the uncovered *positive* correlations.
- Our study demonstrates partial correlation analysis identifies important gene networks underlying growth and susceptibility to the weevil in spruce. In particular, we found transcripts that strongly modify the trade-off between growth and defense, and allow identification of networks more central to the trade-off.

## INTRODUCTION

Coevolved host defenses and herbivores have a long-term evolutionary history (Futuyma & Agrawal, 2009). The resource allocation towards growth on one side, and towards defense and/or reproduction on the other side, is governed by suites of genes and their metabolic pathways. Trade-offs between life-history traits involve the hierarchical allocation of resources to biochemical pathways (Worley *et al*., 2003). Thus, major plant defense theories surrounding a trade-off between growth and defense mainly center on carbon/nutrient balance and growth differentiation balance and the associated allocation costs for defense at the expense of growth and reproduction (Bryant *et al*., 1983; Herms & Mattson, 1992). It is well known that certain signaling pathways usually required for developmental processes (reproductive development, *e.g*., Howe & Jander, 2008; Steppuhn & Baldwin, 2008) can be co-opted for biotic stress responses that directly influence the performance of the pest or contribute indirect defense responses to attract predators or herbivore parasitoids (Thaler *et al*., 2002). This provides the plant with inducible defenses against herbivores or pathogens that are evolutionary advantageous (Steppuhn & Baldwin, 2008). Moreover, when plant growth is decreased by drought for example, then, more carbon becomes available for the production of secondary compounds. Thus, drought stress was shown to increase oleoresin concentration in conifers’ woody tissue with the consequence of higher resistance against herbivory (Turtola *et al*., 2003). Regarding the genetic underpinnings of a growth/defense trade-off, genomic investigations can disentangle this myriad of interactions and demonstrate evolutionary trade-offs between growth and defense. In this paper, we adopt partial correlation analysis to help disentangle such trade-off interactions between growth and defense in trees.

White spruce (*Picea glauca*) and Interior spruce (*P. glauca x engelmannii*) are the economically most important forest tree species in British Columbia. However, both species suffer from infestations by the spruce shoot weevil (*Pissodes strobi*), which consumes the phloem of the apical leader and deforms the main stem (Alfaro *et al*., 2004; vanAkker *et al*., 2004; King *et al*., 1997; He & Alfaro, 2000; Kiss & Yanchuk, 1991; Tomlin *et al*., 1998). Under the pressing need to understand the genetic control of resistance, studies with *Picea* have illustrated host defenses to phloem feeding insects (Ralph *et al*., 2006; Lippert *et al*., 2007; Miller *et al*., 2005; Byun-McKay *et al*., 2006; McKay *et al*., 2003). Ralph et al. (2006) originally documented herbivore-induced transcriptome variation, reflecting either reallocation of resources from primary processes to active defense, or the mobilization of the resources for host tolerance (Howe & Schaller, 2008). Anatomical and the associated chemical defenses in *Picea* bark have also been described (Franceschi *et al*., 2005). The physical structures studied in most detail are the parenchyma cells, and the resin ducts that are all located in the secondary phloem and cambium; the traumatic resin canals are formed in the secondary xylem (Franceschi *et al*., 2005).

In *Picea*, the genetic relationship of insect resistance with height growth is ambiguous (Kiss & Yanchuk, 1991; King *et al*., 1997; Alfaro *et al*., 1997; He & Alfaro, 2000; Lieutier *et al*., 2003). In Sitka spruce, interestingly, genetic resistance was highest in families with only average growth rate (Alfaro *et al*., 2008). Also, constitutive and induced defenses do not always follow sequentially: some resistant trees do not produce/rely on traumatic resinosis (a direct measure for resistance); however, some trees from susceptible families do defend with intensified resin flow to wounding (Alfaro, 1995).

Therefore, in the present study, our specific objective was to identify genes underlying the trade-off between growth rate and weevil resistance. In an experiment initiated by René Alfaro (Alfaro *et al*., 2004), *Picea glauca x engelmannii* progenies were measured for pre-attack plant growth and post-attack damage with several morphological measures, which formed the basis of our hypotheses about correlations. We then estimated gene expression in bark tissue from terminal leaders using microarrays spotted with 13,980 ESTs and their annotated respective transcripts. For network inference, we used a new approach where, with respect to joint values of growth and weevil susceptibility (host defense) traits, we identified a subset of transcripts that exhibited either strong excessive (1) positive partial phenotypic correlations or (2) negative partial phenotypic correlations. That is to say, we chose ESTs that had strong indirect effects on both growth and susceptibility to the weevil, as either jointly positive or jointly negative. These subsets of transcripts were then subject to network analysis, but using entire correlations as the measure of similarity. Our results illustrate how partial correlation analysis of transcript levels can help reveal genes involved in the trade-off between growth and resistance.

## MATERIALS AND METHODS

### Interior spruce pedigree

The experimental population of Interior spruce (*Picea glauca* (Moench) Voss x *P. engelmannii* Parry ex Engelm.) at the Kalamalka Research Station in Vernon, British Columbia (BC), Canada, established in 1995, comprises of 42 full-sib families, each of size 75, from parents selected throughout the Prince George Seed Planning Zone in Northern BC (Alfaro *et al*., 2004). On the basis of previous weevil attack, parents were classified as resistant (R) or susceptible (S), so that families could be grouped as deriving from two resistant parents (R*R: 16 families), one resistant and one susceptible parent (R*S: 20 families), or both susceptible parents (S*S: 6 families). These 42 families were scored for growth and resistance traits. The experimental setup and sampling details are as described in Porth et al. (2011).

### Phenotypic assessments

Initial (early spring) height was recorded in years 1995-1999, terminal leader length was measured in 1999, and weevils were artificially applied in October 1999. After Alfaro *et al*., (2004), attack rates in 2000 and 2001 were classified as (1) successful top kills, (2) attack but no death of leader, and (3) ‘no attack’. The number of eggs laid (oviposition) was also visually recorded as five discrete classes of egg punctures: (1–25), (26–50), (51–75), (76–100), (101+); these were easily distinguished from feeding punctures (R. Alfaro, pers. comm.). The sums of weevil attacks and oviposition for 2000 and 2001 were also used as resistance traits. For the 1999 growing season, bark histology measures were taken for 10 trees per cross, with resin duct measurements taken on upper laterals closest to the leader (Alfaro *et al*., 2004). In the present study we retained only such resin canal characteristics where a positive linear relationship between the leader and the laterals from the same whorl had been shown (Alfaro *et al*., 2004). The following abbreviations for labels and formulas related to bark histology measures were used that followed those provided in Alfaro *et al*., (2004): AREALRC [area of large resin canals (microns squared)], AREASRC [area of small resin canals (microns squared)], TAREARC [total area of resin canals, large plus small (microns squared)], LRC_BA [AREALRC/BARKAREA], SRC_BA[AREASRC/BARKAREA], TRC_BA[TAREARC/BARKAREA], NOLRC[number of large resin canals], NOSRC [number of small resin canals], TOTNORC [total number of resin canals (large plus small)], SZ_IN [AREALRC/NOLRC], NMMS_IN [NOLRC/BAMM], NMMS_OUT [NOSRC/BAMM], NMMS_TOT [TOTNORC/BAMM], BTHK [bark thickness in mm], BARKAREA [total quadrant bark area (square microns)], and BAMM [bark area in mm squared].

### Tissue collection, RNA preparation, microarray, and gene expression profiling

Weevil activity was also observed when we sampled in spring of 2006, and precise records of attacks on-site due to an elevated weevil population were available from 2000-2003. However, for this particular experiment, newly attacked shoots/tree leaders were intentionally excluded from sampling in order to be able to perform an experiment on constitutive levels of defense traits. Thus, we sampled at the right time point, when natural weevil activity occurred in the field. Bark and phloem tissue were collected in the mornings of May 16-18, 2006 and frozen in liquid nitrogen. As we were measuring gene expression, every effort was made to randomize and standardize the collections of tissue. Total RNA was isolated following (Kolosova *et al*., 2004) and quantified via NanoDrop® ND-1000 spectrophotometer; RNA integrity was evaluated using Agilent 2100 Bioanalyzer. The 21,840 spruce ESTs microarray we used for gene expression profiling in this study and the microarray’s quality control are described elsewhere (S. Ralph and co-workers, Gene Expression Omnibus database GEO: GPL5423). A total of 13,980 annotated spruce EST elements were retained for this study. Hybridizations and image acquisition were carried out as described in Ralph et al. (2006) and Porth *et al*., (2011).

### Microarray experimental design and processing of data

A subset of four R*S families, with wide segregation for weevil resistance, were chosen for the gene expression assay. The hybridizations profiled 188 individuals: 48, 36, 50, and 54 in crosses #26, #27, #29, and #32, respectively. After quantitation of the signal intensities in each array, the local background was subtracted for each sub-grid. Data were further normalized using the variance stabilizing normalization method implemented in R package ‘vsn’ (Huber *et al*., 2002). All slides underwent simultaneous normalization to yield a similar overall expression level and variance for each channel independent of the array. A linear model that incorporated dye and block effects in a two-colour microarray design was used (Porth *et al*., 2011). Signal intensities for the original and the normalized measurements were deposited in GEO under GSE22116.

### Partial correlation analyses

Using family means, we estimated genetic correlations in the 42 *P. glauca x engelmannii* crosses. We found: 1) genetic correlations between growth and susceptibility were negative but not strong (*r*∽-0.2, *P*<0.05), 2) genetic correlations between bark histology measures and attack severity were negative and stronger (*r*∽-0.5, *P*<0.05), and 3) genetic correlations between histology and growth were all positive (Fig. S1). These are in accord with previous studies (Kiss & Yeh 1987; Kiss & Yanchuk 1991; King *et al*., 1997) and serve to frame our hypothesis that growth and attack have a negative genetic correlation, but show an overall positive phenotypic correlation (*r*=0.85, *P*<0.05).

To explore the gene networks underlying relations between growth and attack, we used partial correlation analysis to examine how the correlation between two variables (growth and attack) is influenced by a third variable, a gene expression locus (E). As we assayed 13,980 ESTs, we sought to identify ESTs of major effect, and to identify these, most obviously we can inspect bivariate relationships. Fig. 1 plots the observed amounts of growth and attack across 1255 ESTs that had a significant association (*P*<0.01) of expression with the two traits (growth was measured by leader length in yr 1999, and attacks were summed over yrs 2000 and 2001; correlations were recomputed in 1,000 randomized datasets). The phenotypic strength of association almost exactly matches the observed phenotype correlation of 0.85 between growth and attack, but this by itself does not provide any further inferences, especially about networks.

**Figure 1:**
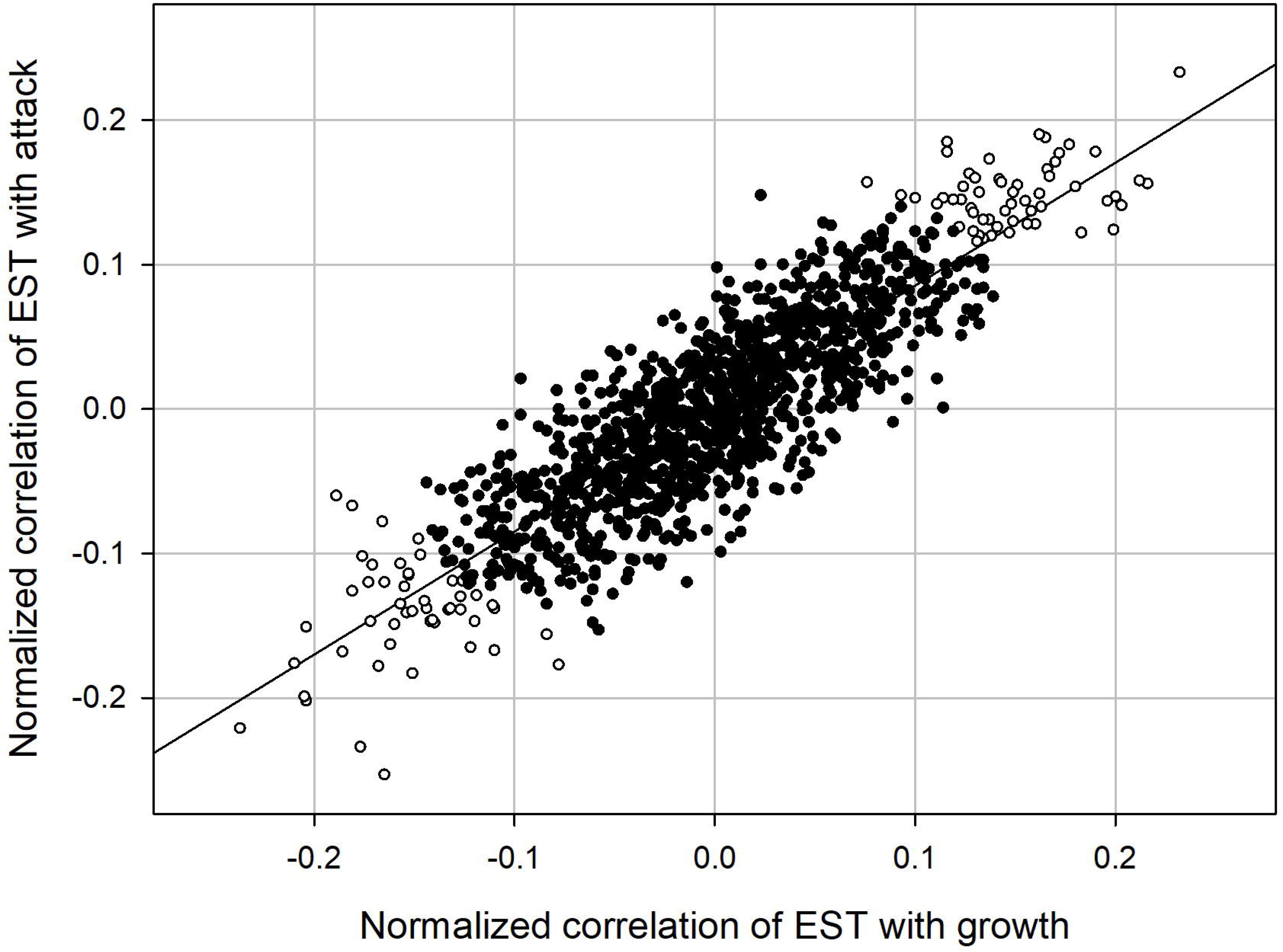
The joint distribution of gene expression in relation to growth (leader length in yr 1999) and susceptibility (number of attacks in yrs 2000 and 2001). Only ESTs that showed associations with a permutation probably of 0.01 or lower are shown. The two ends, which have unshaded circles, illustrate the two groups (*i.e*. the upward bounds in the positive or negative direction of correlations, respectively) that could be subject to separate partial correlation analysis (as demonstrated in the present study).

To this end, we employ the logic of partial correlation to find nodes that are more central to the gene network underlying growth and defense. Let *E* be an expressed gene, *X* be one trait (growth or susceptibility) and *Y* be a second trait (growth or susceptibility). The following are the observed phenotypic correlations: *r_XY_* between *X* and *Y*, *r_XE_* between *X* and *E*, and *r_YE_* between *Y* and *E*. The partial correlation between *X* and *Y*, that is the correlation after the confounding effect of *E* is removed, is

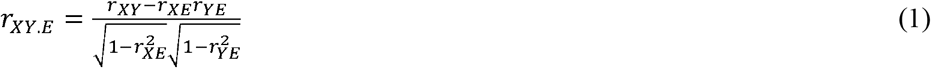

A significant partial correlation can be caused by a mutual effect of *E* on both *X* and *Y*; this can occur when E is at the more central node to *X* and *Y*, relative to other possible nodes. If *r_XE_* and *r_YE_* are either both positive or negative, the partial correlation will be reduced, otherwise the partial correlation will be increased. Regardless of the direction of change, the amount of change is proportional to the relative importance of *E* in the network of *X* and *Y*. A measure of the relative change due to the introduction of *E* is

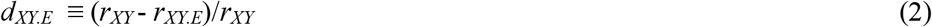

Greater values of *d_XY.E_* occur when *E* explains a greater fraction of the correlation between *X* and *Y*, or generally when *E* is in a tighter network with *X* and *Y*.

We found the more direct connections within a network by systematically evaluating Equation (2) for all pairs of quantitative traits *X* and *Y*, for each EST *E*. Values of *d_XY.E_* greater than a significance threshold are retained in the network; executing this over all possible edges results in a network of putative direct interactions. In constructing the final network, we used total correlations as opposed to the partial correlation quantities in Eq. 2, which only serve to identify ESTs to retain in the network.

A partial correlation threshold network (PCTN) can also be established with rigorous application of a threshold (Kenett *et al*., 2010), but we simply identified two groups. One group of 21 had all positive *d_XY.E_* values and the second group of 166 had all negative *d_XY.E_* values; these groups are described in Notes S1. Significance was again determined by randomization and ESTs of extremely high probability were retained (see Notes S1). One can imagine each group is the average of several tips in Fig. 1 (there are *n*(*n*-1)/2 tips for *n* traits). We note that, in principle, groups of ESTs with all ++ or – *X* vs *Y* effects, and with all +- and -+ *X* vs *Y* effects, could be grouped as co-variates in a multivariate partial correlation analysis.

## RESULTS

### The genetic basis of susceptibility to weevil attack/growth trait correlations (genetic pleiotropy) in P. glauca x engelmannii

Total correlations in the two-by-two factorial R*S spruce progeny are given in Table 1. We identified numerous significant positive and negative correlations between gene expression levels for individual ESTs and the phenotypic traits (Table S1). In addition, *p*-values were computed for each gene-by-growth effect, and then q-values were calculated to adjust for false discovery rate (Storey & Tibshirani, 2003). The effect estimate for each gene was also obtained representing the change in its expression per unit change in growth rate. This identified 867 genes at *q*<0.1, Table S2.

**Table 1:** Number of significant correlations (*P* <0.05; 1,000 randomizations) between gene expression levels testing 13,980 spruce genes and 10 phenotypic traits related to tree height and weevil resistance, respectively, for the two-by-two factorial R*S spruce progeny. Key: ldr99: apical leader length in yr 1999; Init_Ht: initial tree height in yr 1995; Ht_3yrs: tree height in yr 1997; Ht_5yrs: tree height in yr 1999; atk00: attacks in yr 2000; atk01: attacks in yr 2001; atktot: total attacks summed over yrs 2000 and 2001; egg00: oviposition in yr 2000; egg01: oviposition in yr 2001; eggtot: total oviposition summed over yrs 2000 and 2001.

| trait | positive | negative | both | proportion<br>positive |
| --- | --- | --- | --- | --- |
| ldr_99 | 1,827 | 1,553 | 3,380 | 0.540 |
| init_Ht | 844 | 949 | 1,793 | 0.471 |
| Ht_3yrs | 925 | 719 | 1,644 | 0.563 |
| Ht_5yrs | 2,118 | 1,968 | 4,086 | 0.518 |
| atk00 | 933 | 1,021 | 1,954 | 0.477 |
| atk01 | 521 | 630 | 1,151 | 0.453 |
| atktot | 996 | 1,116 | 2,112 | 0.472 |
| egg00 | 640 | 907 | 1,547 | 0.414 |
| egg01 | 474 | 494 | 968 | 0.490 |
| eggtot | 846 | 993 | 1,839 | 0.460 |

We also determined the partial correlations by using the ESTs as independent variables and estimated the relative differences between the total and the partial correlations (Table 2). These pairwise comparisons (*i.e*. the pairwise growth-resistance trait comparisons representing the partial correlation in either direction between growth and susceptibility to weevil attacks) showed large transcriptome responses in terms of the partial correlations representative of the pairwise comparisons between traits (Fig. S2) along with the different effects (*i.e*. the negative and the positive directions, respectively) of the components of these correlations between susceptibility to weevil attacks and growth (Notes S1 for details). Thus, we could uncover all possible combinations related to our initial hypothesis testing regarding the directionality of the genetic correlations to be tested with individual traits (Fig. 1).

**Table 2:** Partial correlations between growth and susceptibility for the two-by-two factorial R*S spruce progeny. Key: ldr99: apical leader length in yr 1999; Ht_5yrs: tree height in yr 1999; atktot: total attacks summed over yrs 2000 and 2001; eggtot: total oviposition summed over yrs 2000 and 2001, in the set of four crosses, grouped by partial correlation analysis where only expression elements with the highest effects on the partial correlation either in the positive direction (+) or negative direction (-) are included in the groups, in groups of size 21, 100 and 166.

| Groups | Trait pairs |  |  |  |  |  |
| --- | --- | --- | --- | --- | --- | --- |
|  | ldr_99 x | ldr_99 x | ldr_99 x | Ht_5yrs x | Ht_5yrs x | atktot x |
|  | Ht_5yrs | atktot | eggtot | atktot | eggtot | eggtot |
| All | 0.858 | 0.287 | 0.339 | 0.265 | 0.297 | 0.815 |
| Top100+ | 0.845 | 0.121 | 0.185 | 0.029 | 0.138 | 0.777 |
| Top100- | 0.821 | 0.448 | 0.570 | 0.435 | 0.514 | 0.792 |
| Tight21+ | 0.813 | 0.118 | 0.229 | 0.075 | 0.168 | 0.785 |
| Tight166- | 0.743 | 0.075 | 0.211 | 0.215 | 0.128 | 0.568 |

We estimated the correlations’ upward bounds involving the top 100 genes in the positive and the negative correlations between the susceptibility to weevil attacks and growth, respectively (Table 2). This provided a clear picture regarding a significant increase (top 100) of the partial correlations coefficient compared to the phenotypic (total) correlations which had shown a positive trend for growth and weevil attack (Table 2). Therefore, the results relating to the partial correlations also indicated that we could uncover the genetic effect of the transcriptome attributed to the negative genetic correlations between weevil attack and height growth in spruce (see above). Still, we found transcripts that could enhance growth but could also have a negative impact on resistance or vice versa (positive correlations), Table 2.

### Identification of candidate networks using partial correlations

The Supplement material (Notes S1) gives the relative partial correlations (Eq. 2) for individual ESTs and each of the six pairwise combinations of the four traits. These four traits were ldr_99 (leader length in 1999), Ht_5yrs (height at age 5) and atktot (attack damage, total), eggtot (egg number, total). There were many significant positive and negative partial correlations between gene expression levels for individual ESTs and the phenotypic traits. Hence, there was sufficient power to use partial correlation analysis to detect networks of interest. Thus, most importantly, on the basis of the partial correlation analysis and Eq. 2, we identified two distinct compact networks (*P*=0.004 for all pairwise comparisons) for the positive correlations (++ or -- effects) involving 21 transcripts (represented by Fig. 2A) and for the negative correlations involving 166 transcripts (corresponding to +- or -+ effects), respectively (represented by Fig. 2B), and in Fig. 1, respectively. The details of the genes’ identities for these two networks are provided in Table S3 (corresponding to the network presented in Fig. 2A) and in Table S4 (Fig. 2B). The complete lists of genes used in four transcript files (of sizes 100, 166 and 21, respectively) are given in Notes S1 as well as in Tables S3-S6.

**Figure 2:**
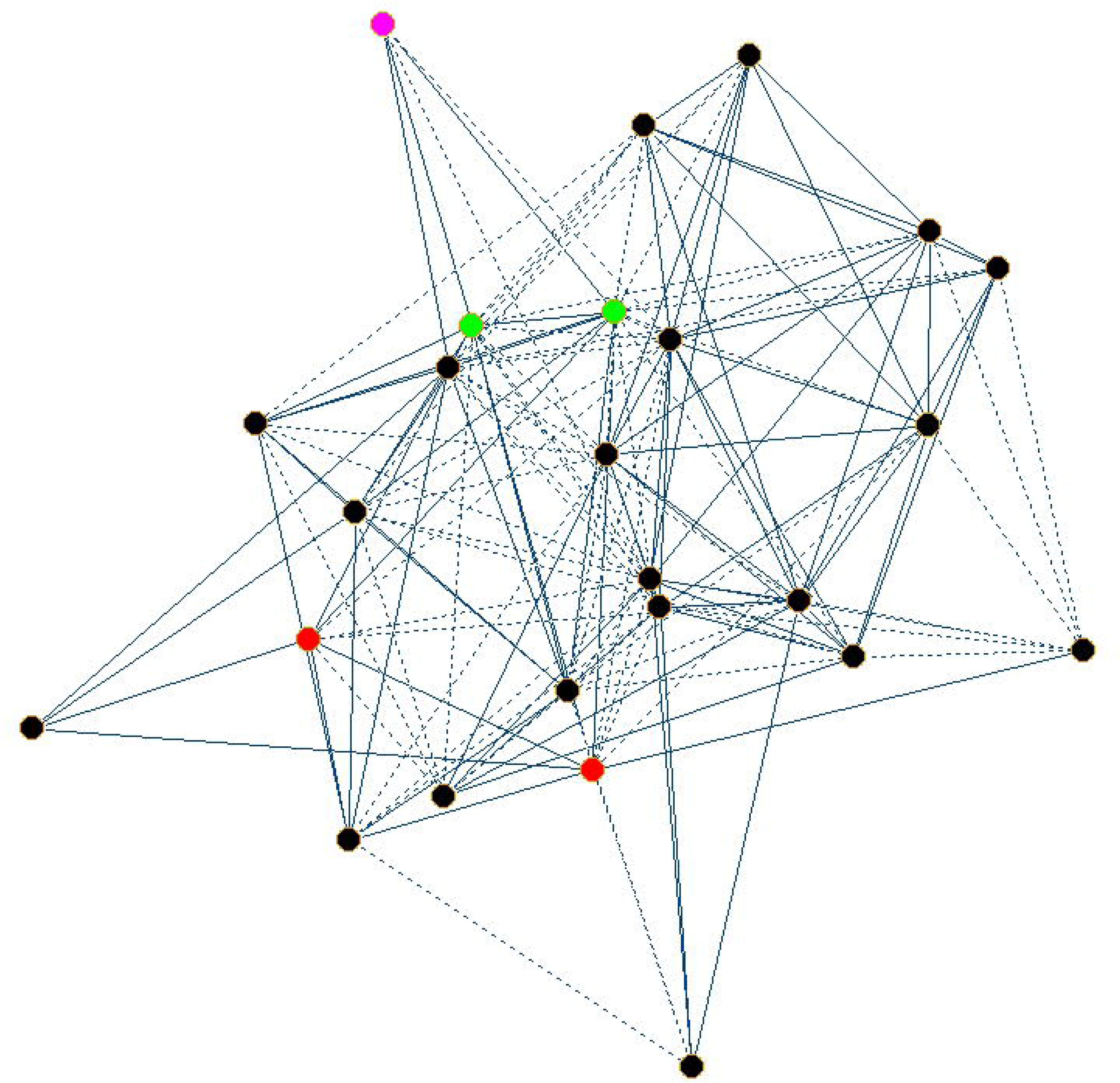
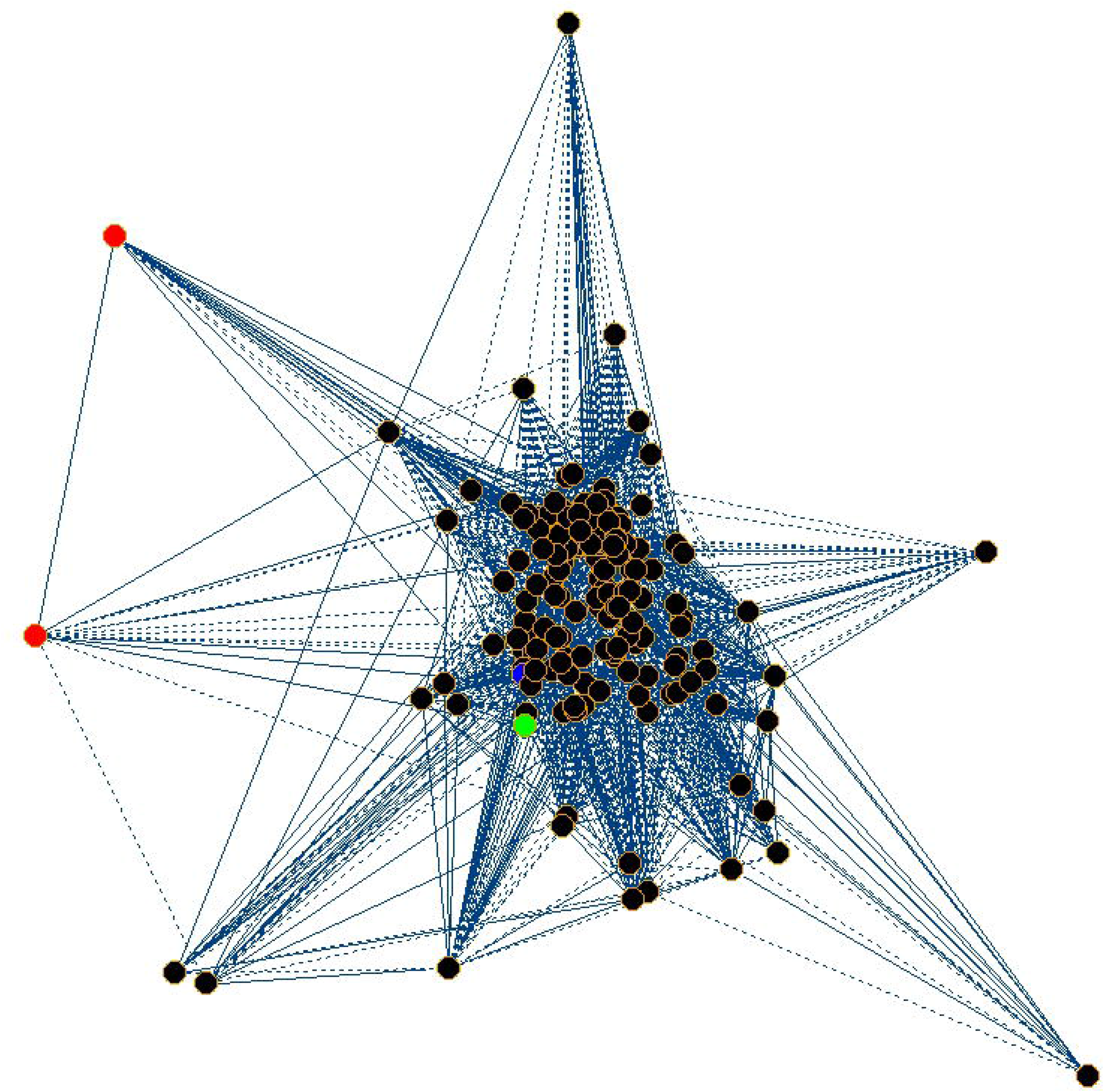
Trade-off vs No trade-off scenarios testing between spruce tree growth and host’s weevil pest resistance for the two-by-two factorial R*S spruce progeny uncovered by correlating the expression of 13,980 (annotated) spruce genes in bark tissue of the apical shoot with host tree height growth and host resistance towards the weevil phenotypes. In both networks, correlation coefficient cutoffs of 0.197 were employed (Notes S2 and S3 show entire results); the social networks were visualized using Pajek (vlado.fmf.uni-lj.si/pub/networks/pajek/) employing the Fruchterman-Reingold approach; dashed lines represent *negative* correlations, solid lines represent *positive* correlations. **A**, Correlation network representative of the *positive* genetic correlations between host tree growth and pest attack among individual traits involving height at age 5 (yr 1999), apical leader length in yr 1999 (green vertices), total attacks and a total number of egg plugs (yrs 2000-2001) (red vertices) and 21 gene transcripts (*P*=0.004) (black vertices, except the spruce *acrocona* ortholog is shown in magenta). Spruce element annotations are provided in Table S3. **B**, Correlation network representative of the *negative* genetic correlations between host tree growth and pest attack among individual traits involving height at age 5 (yr 1999), apical leader length in yr 1999 (green vertices), total attacks and a total number of egg plugs (yrs 2000-2001) (red vertices) and 166 gene transcripts (*P*=0.004) (black vertices, except the spruce *PDR12* ortholog is shown in blue). Spruce element annotations are provided in Table S4.

### The genes at the centre of pleiotropy between tree growth and pest resistance in P. glauca x engelmannii

A putative PDR type ABC transporter *ATPDR12/PDR12 (PLEIOTROPIC DRUG RESISTANCE 12)*, represented as element WS0269_K02 on the 21.8k spruce EST array, showed the strongest significance among all 13,980 tested transcripts in the correlations between gene expression, tree height and weevil attack phenotypes (*P*= 0.019), Table 3. Variation in *PDR12* steady-state gene expression at the population level represents an example of transcripts that were positively correlated with height but negatively correlated with weevil susceptibility traits (+- effects) in the two-by-two factorial crosses progeny widely segregating for resistance. A set of additional genes showed the same directionality in gene expression with growth and resistance variation for this large progeny of 188 individuals originating from 4 different crosses (Table S1).

**Table 3:**
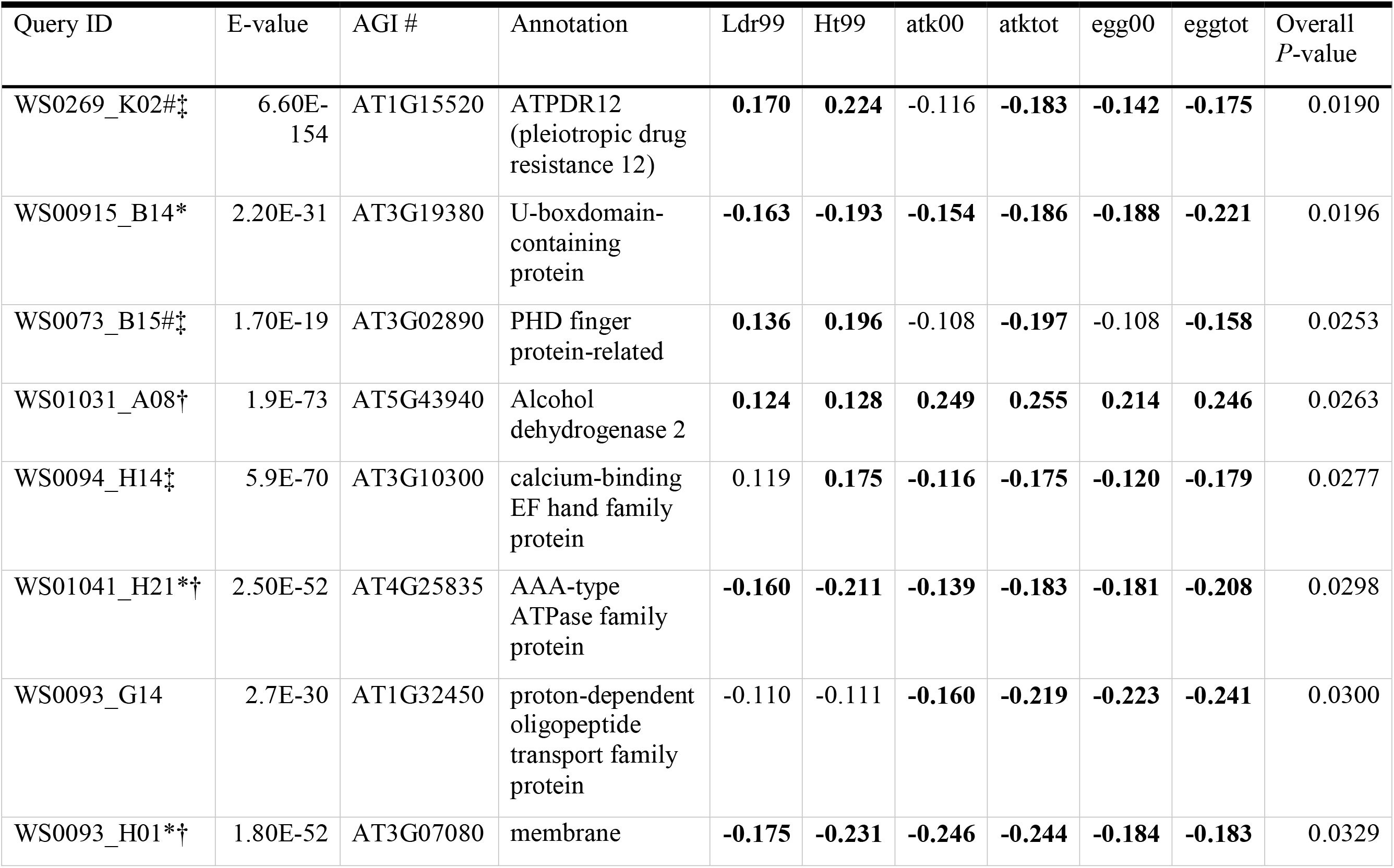
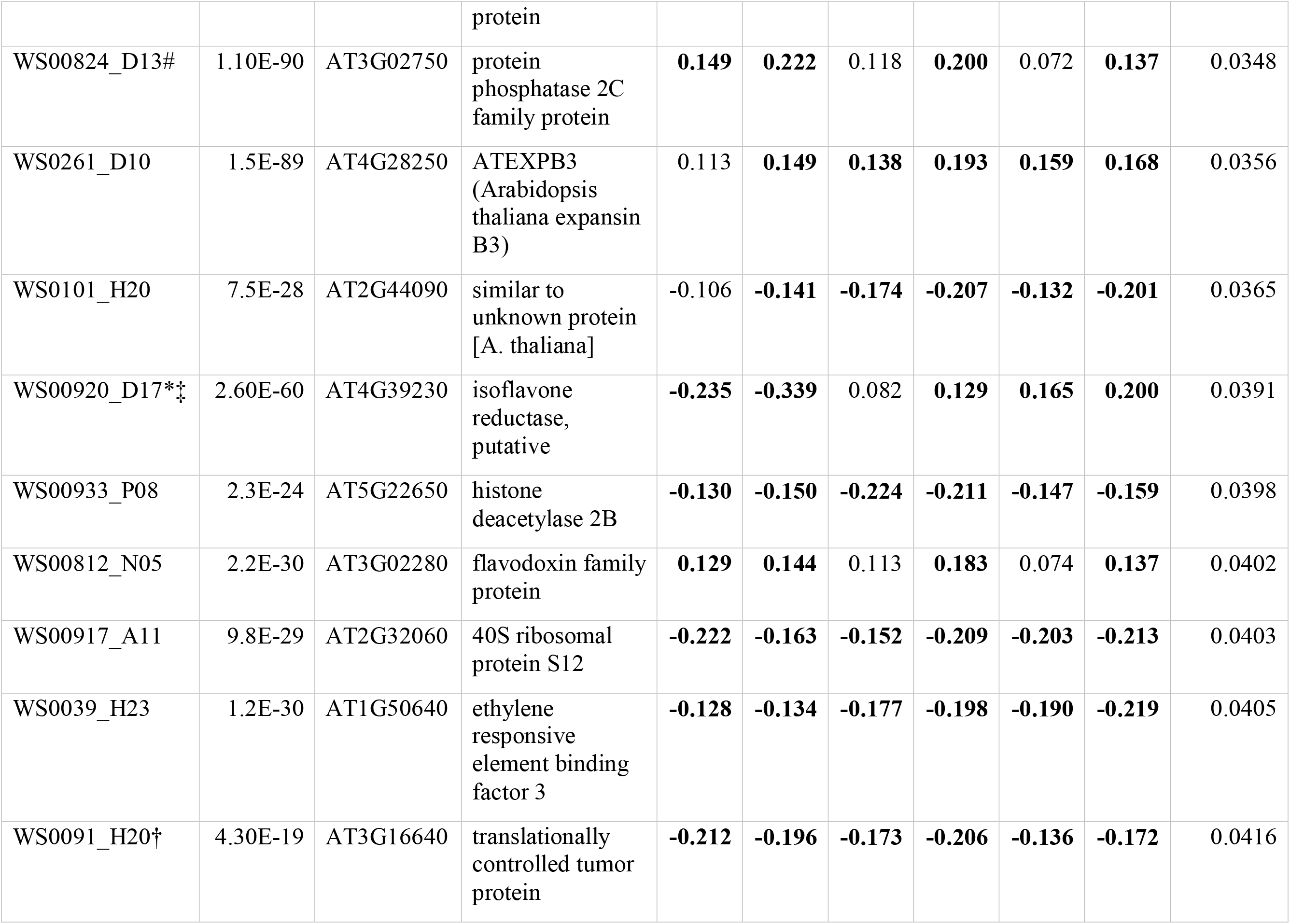
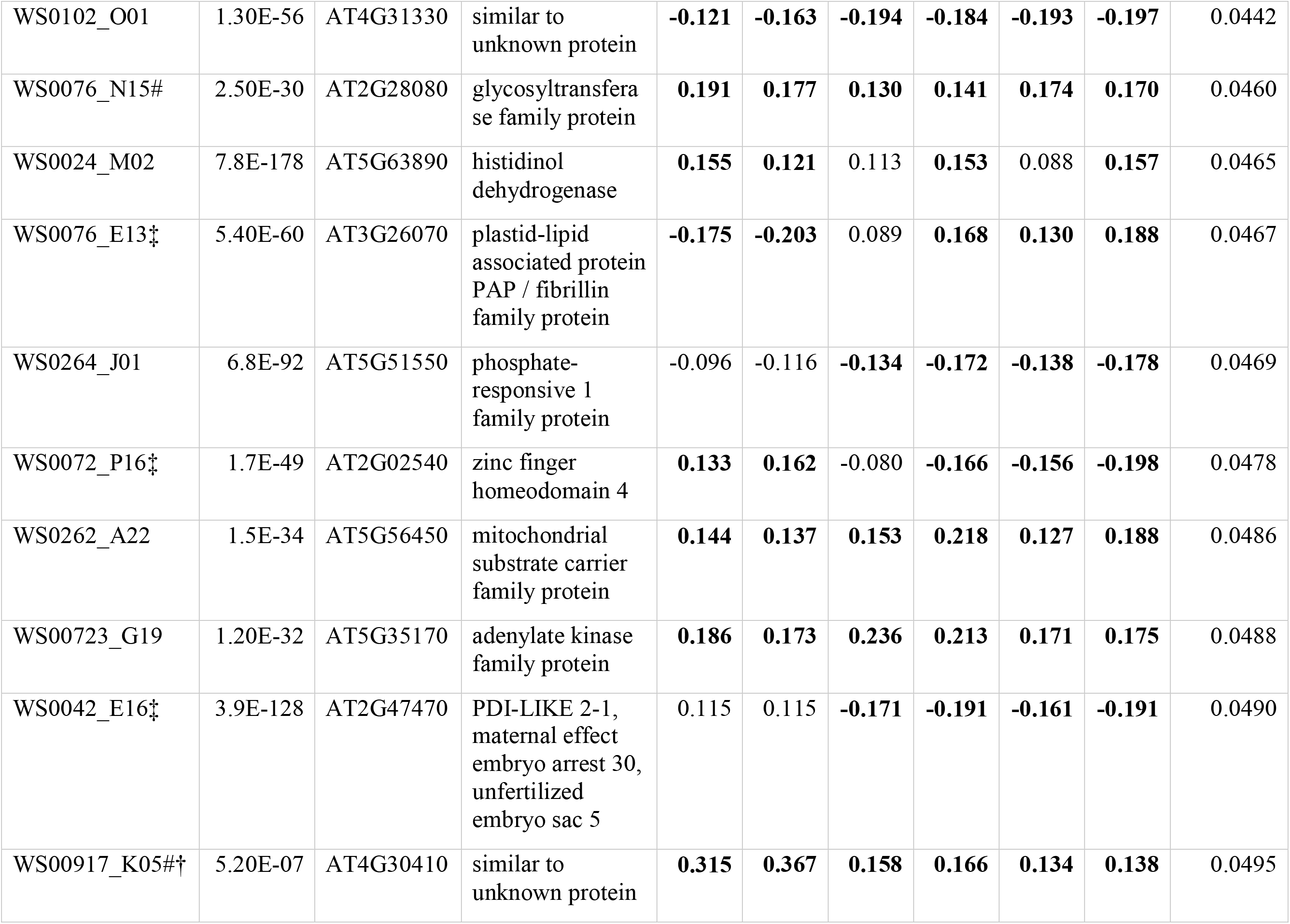
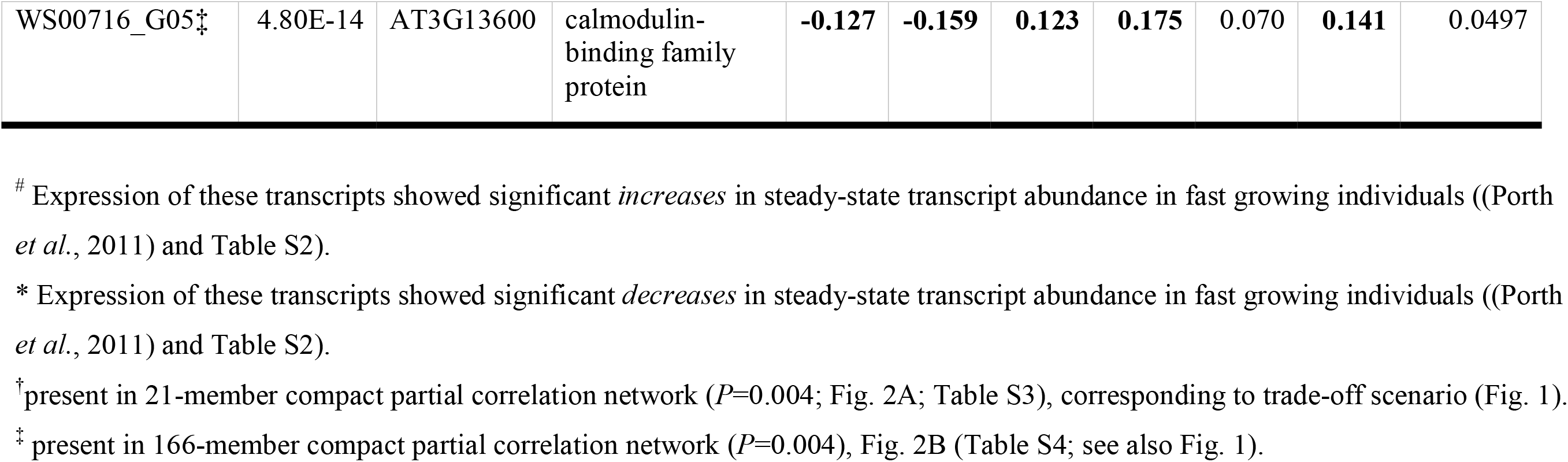
Twenty-eight most significant transcripts from gene expression correlations for the two-by-two factorial R*S spruce progeny with Ldr99, Ht99, atk00, atktot, egg00, eggtot phenotypes (*P*-value < 0.05 in bold; 1,000 permutations), sorted by overall significance Key: Ldr99: apical leader length in yr 1999; Ht99: height at age 5 (yr 1999), atk00: attacks in yr 2000; atk01: attacks in yr 2000, and atktot: total attacks; egg00: oviposition in yr 2000; egg01: oviposition in yr 2001, and eggtot: total number of egg plugs (yrs 2000-2001); sequence homology of the spruce cDNA to the respective Genbank entry (BLAST identity, AGI# for AT homolog) is supported by the expect value (E-value) of the hit.

However, among transcripts that correlated negatively (-- effects) with both growth and attack/oviposition (again tested by 1,000 permutations), there were spruce genes with the following functions: (a) dirigent proteins related to constitutive defenses (array elements WS0086_I04 and WS0104_A04 annotated as *PicsiDIR29* and *PicsiDIR35*, respectively (Ralph *et al*., 2007; Porth *et al*., 2011)), (b) peroxidases important in the reinforcement of anatomical structures through lignification (WS01029_G23, WS01033_K22, and WS01017_F04 (Porth *et al*., 2011)), (c) other cell wall modifying proteins such as xyloglucantransglycosylases (WS0264_P12, WS0041_D16), xyloglycosyltransferases (WS0072_C14, WS00918_N14), pectinesterase (WS0039_I15), (d) stress inducible peroxidase *PicabPIPRX*(WS01029_D16 (Porth *et al*., 2011)), stress signaling pathway related AP2 domain transcription factor *TINY* (WS0102_C24) (Sun *et al*., 2008), (e) flavonoid-based defense related *F3’H* (WS00931_D17 (Porth *et al*., 2011)), and finally (f) the weevil-inducible terpene synthase (WS0078_K20 (Kolosova, 2010)). Three candidates that showed the negative correlation with growth, the chaperonin heat shock protein 60 WS00920_E06, the ethylene inducible universal stress protein family protein WS01027_A08, as well as the terpene synthase WS00929_B22 were found to be significantly downregulated in weevil-resistant spruce trees (Verne *et al*., 2011).

Our results indicated that expression of these three genes negatively correlated with attack rates, and their expression was also negatively correlated with growth; therefore we assume a potential trade-off. Among all these genes mentioned above, only three were represented in the 21-member gene-gene interaction network (the two spruce peroxidases potentially implicated in lignification, WS01033_K22, and WS01017_F04, as well as the spruce AP2 domain transcription factor *TINY* WS0102_C24), Fig. 2A and Table S3.

## DISCUSSION

Our work investigated the co-evolution of height growth vigour and pest resistance in spruce and employed transcriptomics data to obtain insights into this relationship. We assayed for gene expression levels in a sample of four families (188 individuals in total) segregating for weevil resistance. Gene expression levels were interrogated for 13,980 annotated ESTs spotted on a 21.8k member microarray. We then estimated the partial correlations of pairs of growth/attack traits with gene expression levels, and identified ESTs of high correlation likely at the centre of such phenotypic trait correlations among growth and attack traits.

In conifers, the generalized strategy for protection against bark boring pests are the constitutive defenses that are localized within the periderm, the cortex, the secondary phloem and xylem and are arranged in concentric, multiple layers (Franceschi *et al*., 2005). Toxins, antifeedants, defensive proteins and enzymes, and reservoirs of chemicals such as resins are released from the bark upon attack. The spruce shoot weevil attacks the host tree at the shoot apical leader of previous year’s growth (Kiss & Yanchuk, 1991). The apical leader unites the high fitness value for the tree’s competitiveness with high probability for weevil attack; consequently, this part of the tree receives priority in the allocation of defense metabolites. The attractiveness of the prospective host to the herbivore is highly related to the host’s nutritional adequacy (*e.g*. content of sterols, lipids, fatty acids, amino acids, carbohydrates). Since preformed defenses are established coordinately during the development of secondary xylem in the apical shoot (Friedmann *et al*., 2007), the tree’s innate metabolism related to growth and normal development very likely influences the establishment of these defenses directly, thus forming the foundation for genetic pleiotropy. In the present study, we further explored the evidence of pleiotropy between growth and constitutive resistance against the herbivore in the host (Porth *et al*., 2012) including a potential trade-offs.

We investigated the correlations between growth and resistance traits involving transcript expression to identify genes that might be at the centre of potential trade-offs between inherent growth rate and constitutive defenses. Transcriptomics can uncover causality in true interactions between genes, if in the analysis of pairwise gene interactions all other transcripts are also considered (partial correlation analysis, c.f. Johansson *et al*., 2011). Consequently, our correlation networks based on such analysis highlighted the more direct associations among the myriad of all possible associations.

### JA biosynthesis genes

Hormonal crosstalk is indispensable for regulating the growth-defense trade-offs and jasmonate (JA) signaling lies at the centre of networks that define defense strategies against herbivory (Huot *et al*., 2014). The differential regulation of certain components/steps in the JA pathway generates distinct responses to different stimuli (reproductive development, growth or types of defenses that can be active defenses (Kazan & Manners, 2008)). Following the apparency theory, fast growing individuals are thought to be biased towards induced defenses (Steppuhn & Baldwin, 2008). The chemical compounds providing active (induced) protective defenses against herbivory are the terpenoids and some phenolic derivates (such as tannins (Bauce *et al*., 2006)). In general, the induced defenses rather acquire signaling systems from developmental programs; and such defenses are expressed only when required. Hence, such types of defenses primarily allow resource investment for the plant’s competitiveness through increased growth (and reproduction). For example, one study failed to connect defensive tannin production with a growth trade-off (Haering *et al*., 2008), but see further below. In our study, the levels of JA defense signaling in fast growing (unattacked) progeny were lowered. Genes involved in the plastid located early JA biosynthetic steps (*LOX1:* WS01014_A24; *AOS CYP74A:* WS01016_F05 and *AOC2* genes: WS00820_E17) all showed decreases in steady-state transcript abundance in fast growers (Table S2). We also retrieved these genes in the compact 166-member network involving the negative correlations between growth and attack (Fig. 2B). Also, the majority (c.70%) of the suite of the spruce genes involved in reproductive (male and female strobili) development, identified to be differentially expressed, exhibited increases in steady-state transcript abundance in fast growing (unattacked) progeny (Table S2).

### Terpenoid-based defense

We identified two spruce genes with roles in terpenoid defense against herbivory in the compact 166-member network (Fig. 2B). First, we found *PDR12*, which is possibly involved both in constitutive as well as in induced defense. In *Arabidopsis, PDR12* shows a response to infection by necrotrophic fungal pathogens but its upregulated gene expression can also be artificially elicited by applications of methyljasmonate, salicylic acid and ethylene in growth media; its study revealed the first evidence of *active* transport of terpenoids in plant defenses (Campbell *et al*., 2003; van den Brule & Smart, 2002). Second, we also identified a putative linalool oxygenase (*CYP76C1*) with a potential role in detoxification during plant-insect interactions (Hofer *et al*., 2014). Interestingly, expression levels of these two genes showed marked differences with growth rate, with *PDR12* increasing in steady-state transcript abundance in fast growers. Other spruce genes implicated in terpenoid metabolism including genes from the methylerythritol phosphate pathway, monoterpene (myrcene, pinene), sesquiterpene (farnesene), or tocopherol biosynthesis exclusively showed increases in steady-state transcript abundance in fast growers. Tocopherols promote stress tolerance by protecting against oxidative stress, while terpenoids have mostly active defensive functions. Interestingly, these genes were not identified in previous weevil feeding experiments (where a very limited number of tested genotypes was included), thus they were also not functionally characterized previously (Ralph *et al*., 2006; Keeling *et al*., 2011). Possibly, the phenotypic expression of tolerance and well-established chemical defenses towards herbivory is mutually exclusive.

### Flavonoid-based defense

Several flavonoid pathway-related genes were part of the 166-member network (Fig. 2B; Table S4). Functions were related to the biosynthesis of anthocyanins/tannins (protective function against herbivorous insect damage and/or associated fungi (Bauce *et al*., 2006; Hammerbacher *et al*., 2014) and biosynthesis of lignans (via phenylcoumaran benzylic ether reductases, *PBR* (Macrae & Towers, 1984)). And while *PBR PicglPPR05* was associated with attack rates, there was also evidence that this gene was tightly regulated by a secondary growth related transcription factor (*myb20*), and showed decreases in steady-state transcript abundance with increased growth rates (Porth *et al*., 2011), interestingly, with a positive effect on genetic resistance (Table S4). *CYP750* members were related to weevil resistance (Porth *et al*., 2011) but also to growth traits with differentially correlated gene expression. Although such conifer-specific P450 members are supposedly involved in a wide range of derivatization reactions in phenylpropanoid metabolism, their exact functions need yet to be ascertained. Dedicated tannin related and downstream regulated genes showed either no or a positive relationship with growth rate (Table S2) suggesting similar findings as a previous study that failed to relate tannin production to a trade-off (Haering *et al*., 2008). This pattern was opposite to the above mentioned early flavonoid pathway gene *CYP75/F3’H* expression implicated in a potential trade-off. Possible reasons for such trade-off are the cross-talk between the lignin and the flavonoid pathways (metabolic plasticity (Mouradov & Spangenberg, 2014)) and the additional role flavonoids play in plant development (auxin transport: Taylor & Grotewold, 2005). In addition, other studies exist, which showed trade-offs based on phenylpropanoids and condensed tannins, particularly under stress conditions that would change plants’ allocation choices (*e.g*. pine trees: Sampedro *et al*., 2011; Salicaceous trees: McKown *et al*., 2014). Because of such interrelations, further investigations are warranted that also test for plants’ growth-defense relationships under different stress conditions, including such comparisons for constitutive vs induced chemical defenses (see also below).

### Reproductive Development

We identified spruce MADS-box genes that are candidates for reproductive maturity or reproductive meristem identity in conifers [annotated as *AGL20 (Suppressor of Overexpression of Constans 1), SOC1-like* gene *AGL42*, and *AGL2 (SEPALLATA1)* based on sequence homology with *Arabidopsis* genes] (Katahata *et al*., 2014; Uddenberg *et al*., 2013; Melzer *et al*., 2010) which are paralogs of flowering genes in *Arabidopsis*. As there are no *SEP1* genes in gymnosperms, the identified *P. glauca* gene WS00823_F11 is likely the ortholog of *P. abies DEFICIENS AGAMOUS-LIKE1* (PgMADS10_DAL1, with proposed function in regulating the transition from juvenile to adult phase in *Picea* (Carlsbecker *et al*., 2004)), and which is located within the *SEP1* sister clade more closely related to *AGL6*. Likewise, *SOC1* and *SOC1-like* annotated *P. glauca* genes, WS0056_A03 and WS00922_C06, respectively, are homologs of *P. abies DAL3* (PaMADS7_DAL3 (Melzer *et al*., 2010)). Functional analysis of *CjMADS14*, the *Cryptomeria japonica* ortholog of *P. abies DAL1*, and *CjMADS15*, the *C. japonica* ortholog of *P. abies DAL3*, indicated that expression of *DAL1* is specific to the reproductive organs (and its function is related to suppressing reproductive repressors), while expression of *DAL3* is more ubiquitous but included male and female strobili (Katahata *et al*., 2014; Niu *et al*., 2016).

In both the negative and the positive correlation networks, we found the *P. glauca DAL3* homolog WS00922_C06, as well as the *P. glauca DAL1* gene and the other *P. glauca DAL3* homolog, WS0056_A03. Expression of the *P. glauca DAL3* homolog WS0056_A03 showed significant increases in steady-state transcript abundance in fast growing spruce individuals (*q*<0.05) (conversely to *DAL1* whose expression pattern was unaffected in either direction by growth) and its expression was associated with the identified growth-resistance trade-off, while the second *DAL3* homolog was significantly down-regulated in fast growers (*q*<0.05) and part of the negative correlation between growth and attack (Fig. 2B). These results are indicative of opposing functions of the two *DAL3* homologs and a striking example of subfunctionalization in the extensively expanded and duplicated clade of TM3-like genes in *Pinaceae* (Gramzow *et al*., 2014). Interestingly, *DAL3* is closely related to the *P. abies acrocona (acr42124_1/DAL19*) gene, which promotes early cone-setting (Uddenberg *et al*., 2013), and to which WS0056_A03 would be the closest of the two white spruce homologs. It is important to mention here that similar to the *P. abies acrocona* phenotype (Uddenberg *et al*., 2013), an enhanced WS0056_A03 expression is also negatively correlated with expression of genes involved in cell wall modification, cell signaling, and plant stress response (Fig. 2A and Table S3). Such gene expression pattern was identified as a trade-off between enhanced growth and weevil resistance.

### Different genomic introgression patterns in Interior spruce

We would like to highlight the putatively different genomic introgression patterns among the studied *P. glauca* x *P. engelmannii* hybrid progeny as one possibility to explain the differences in the relationship between growth and resistance among individuals, as, for example, it is known that pure *P. glauca* grows at different rates than pure *P. engelmannii* or their respective hybrids, and this relationship also differs for different life-history stages; while *P. glauca* initially grows more slowly, it outpaces *P. engelmannii* after tree age 10 (De La Torre *et al*., 2014). Current tree breeding programs select towards more vigorous trees thus higher *P. glauca* ancestry, and concurrently restrict the downward displacement of Engelmann spruce seedlots due to increased weevil susceptibility. We point out that there may be differences in selection pressure on the pure species, as low elevation populations are much more resistant to weevils than their high elevation counterparts (which mainly consist of *P. engelmannii*), where weevils cannot thrive. However, there is currently no scientific evidence for significant differences in weevil resistance between the pure species (B. Jaquish, personal obs.).

### Conclusions and prospects

Research on the relationships between plant growth and defenses against herbivory is timely, especially, when physiological with molecular approaches are combined to predict trade-offs (Züst & Agraval, 2017). Thus, for functional genomics studies of host defense mechanisms deployed against herbivores, it is important to consider inherent growth characteristics of the host. This research has not received enough attention in forest trees, although tree breeding strategies seek to optimize the growth-defense balance, and ideally maximize for both growth and defense. Such obtained knowledge is particularly important for conifer trees species, which are widely planted and are already undergoing advanced generation breeding programs, in contrast to deciduous tree species, such as poplars.

Here, we used partial correlation analysis to identify the key genes and network nodes based on gene expression associated with phenotypic and genetic variability in the context of tree resistance to a major pest. Overall, our study did not show a trade-off between high growth rates and defenses. Nevertheless, important environmental components could influence the relationship between growth and pest resistance. Previously assessed phenotypic correlations that were found to be largely positive between shoot growth and susceptibility against weevil attacks would indeed hint at underlying environmental components. To further untangle the gene regulatory networks underlying the conifer tree’s life history strategy, we are now investigating *ABCG40/PDR12* gene expression after wounding, under the alternative conditions of (1) drought stress, and (2) no drought stress imposed. It has been postulated that drought stressed conifers can rely more on constitutive than on induced defenses. If this candidate gene is involved with the trade-off, we can further characterize the role of *ABCG40* in relation to stress, and perhaps, separate the role of constitutive versus induced responses. We can also immediately apply such knowledge to breeding efforts against the stem boring pest, particularly for known local stress conditions, such as drought.

## ACKNOWLEDGEMENTS

This work was supported by Genome BC, Genome Canada, and the Province of BC (Conifer Forest Health Genomics grant to K.R.). We acknowledge Dr. Carol Ritland for project management. We thank the technical staff of the Ritland lab. We further thank Dr. René Alfaro (Canadian Forest Service) for phenotypic data provision.

## AUTHORS’ CONTRIBUTIONS

I.P., B.J. and K.R. performed experiments, conducted fieldwork, and analyzed data. R.W. provided statistical assistance. I.P. and K.R. drafted the manuscript. K.R. and I.P. planned and designed the research.

## SUPPORTING INFORMATION

**Figure S1**: Correlation structure for significant genetic correlations among 18 individual host tree characteristics.

**Figure S2**: Transcriptome changes for the two-by-two factorial R*S spruce progeny in terms of partial correlations.

**Table S1**: Full list of correlation results of gene expression for the two-by-two factorial R*S spruce progeny and for the 10 quantitative traits.

**Table S2**: Effect of inherent growth rate variation on global transcriptome changes for the two-by-two factorial R*S spruce progeny.

**Table S3**: Gene id’s of the 21-member compact network (*P*=0.004).

**Table S4**: Gene id’s of the 166-member compact network (*P*=0.004).

**Table S5**: Top100+ transcripts, among all four pairwise growth-resistance trait comparisons.

**Table S6**: Top100- transcripts, among all four pairwise growth-resistance trait comparisons.

**Notes S1**: Partial correlations for the two-by-two factorial R*S spruce progeny.

**Notes S2**: Correlation network 21+, full results.

**Notes S3**: Correlation network 166-, full results.

